# DNA Base Pair Polarities of Pyrimidine(s)-Purine(s) (Pyu) dsDNA induce stronger Hybridization and unique DNA Conformation compared to Purine(s)-Pyrimidine(s) (Puy) dsDNA

**DOI:** 10.1101/2024.08.23.609369

**Authors:** Suk Keun Lee, Dae Gwan Lee, Yeon Sook Kim

## Abstract

The present study introduces a concept of DNA base pair polarity in double-stranded DNA (dsDNA), which has led the serial investigation of hydrogen bonding magnetic resonance-based gene regulation. DNA base pair polarity is characterized by hydrogen bonding between electrostatically positive purine and negative pyrimidine, exerting the stabilization and organization of dsDNA. Comparison of DNA hybridization potential between Pyu (pyrimidine(s)-purine(s)) and Puy (purine(s)-pyrimidine(s)) oligo-dsDNAs in EtBr intercalated electrophoresis showed that Pyu ds(nTnA) (n=3, 4,5,6,7,8,10,12) and ds(nCnA) (n=7,8,10,12) exhibit stronger hybridization than Puy ds(nAnT) and ds(nAnC), respectively. In the HPLC for DNA conformation change in 0.1M NaCl solution, Pyu oligo-dsDNA tended to have a larger peak area than Puy oligo-dsDNA, indicating that the former had a more extended and organized DNA conformation than the latter. FT-IR results showed that Pyu ds3T3A, ds3C3G, dsCTCGAG, and dsTCTAGA showed higher IR absorbance at 3400-3200 cm^-1^ than Puy ds3A3T, ds3G3C, dsGAGCTC, and dsAGATCT, respectively, indicating Pyu oligo-dsDNAs have increased N-H stretching in hybridization than Puy oligo-dsDNAs. *In vitro* RNA transcription from template plasmid DNAs inserted with six segments of Pyu ds6(TCTGA) and positive ds14(GA) produced more RNA than those with seven segments of palindromic Pyu ds(CATG) and negative ds14(CT), respectively, indicating the former plays a stimulating role for RNA transcription while the latter plays an inhibitory role. When comparing PCR primers between Pyu ds12/n(nCnA) and Puy ds12/n(nAnC) (n=1,2,3,4,6,12), Pyu primers can easily hybridize on template DNA and subsequently produce abundant DNA compared to Puy primers. The data suggest that Pyu oligo-dsDNAs hybridize strongly and form distinctive conformation, thereby stimulating RNA transcription more than Puy oligo-dsDNAs. Since Pyu oligo-dsDNA appears to preserve DNA signal and conformation relevant to DNA function more efficiently than Puy dsDNA, it is proposed that the entire DNA code can be divided into Pyu dsDNAs with unique DNA base pair polarities.

---

The structure of dsDNA is fixed by A-T and G-C pairing and has a large sequence diversity resulting in 4^n^ (n=number of base pairs (bps) in dsDNA) different dsDNAs, however, the sequence-specific DNA signal could be illustrated simply by electrical polarities. A computational study of different DNA topologies in electrolyte solution also revealed divalent counterions on the effective pair potentials (*1*). This study provides a molecular interpretation of DNA base pair polarities, which represent the electrostatic potential of dsDNA and may have an impact on the regulation of DNA function.

## DNA base pair polarities encoding Pyu and Puy dsDNAs

The dsDNA is known to contain genetic information to interact with specific molecular polarities and conformations of transcription factors. It is composed of complementary base pairs hybridized by hydrogen bonding between positive purine and negative pyrimidine. A strand of dsDNA has a functional direction from 5’ to 3’, which is determined by the intramolecular polarity from negative phosphate group to positive base in a nucleotide. Although DNA sequence looks random distribution of nucleotides, A, C, G and T, the linear sequence of nucleotides is characterized by their molecular polarities.

With an additional imidazole ring to pyrimidine ring, purine may exert stronger intramolecular electron resonance than pyrimidine (*2*). This may facilitate the intramolecular electrostatic shift from pyrimidine to purine while impeding from purine to pyrimidine, resulting in the semi-conductive property in a single strand of dsDNA (*3*). Therefore, there are two distinct electrostatic shifts in the nucleotide sequence, an electrostatically conductive from pyrimidine to purine, Pyu sequence, and an electrostatically resistive from purine to pyrimidine, Puy sequence. And then, the Pyu sequence can facilitate the intramolecular electrostatic shift to its base rings via electron resonance more than the Puy sequence.

This concept allows clear visualization of potential electrostatic interactions between adjacent base pairs, as well as the boundary between pyrimidine(s) and purine(s) in forward and reverse DNA strands. The electrostatic shift from pyrimidine(s) to purine(s) is enhanced by the forward attraction of the purine-dependent phosphate group, while the electrostatic shift from purine(s) to pyrimidine(s) is hindered by the reverse attraction of purine-dependent phosphate group of a DNA axis. Therefore, we hypothetically assume the existence of an electrostatic shift boundary between purine(s) and pyrimidine(s) in a DNA strand and draw a boundary line mark between purine(s) and pyrimidine(s) in the symbolic DNA graph (Fig.1).

**Fig. 1.**
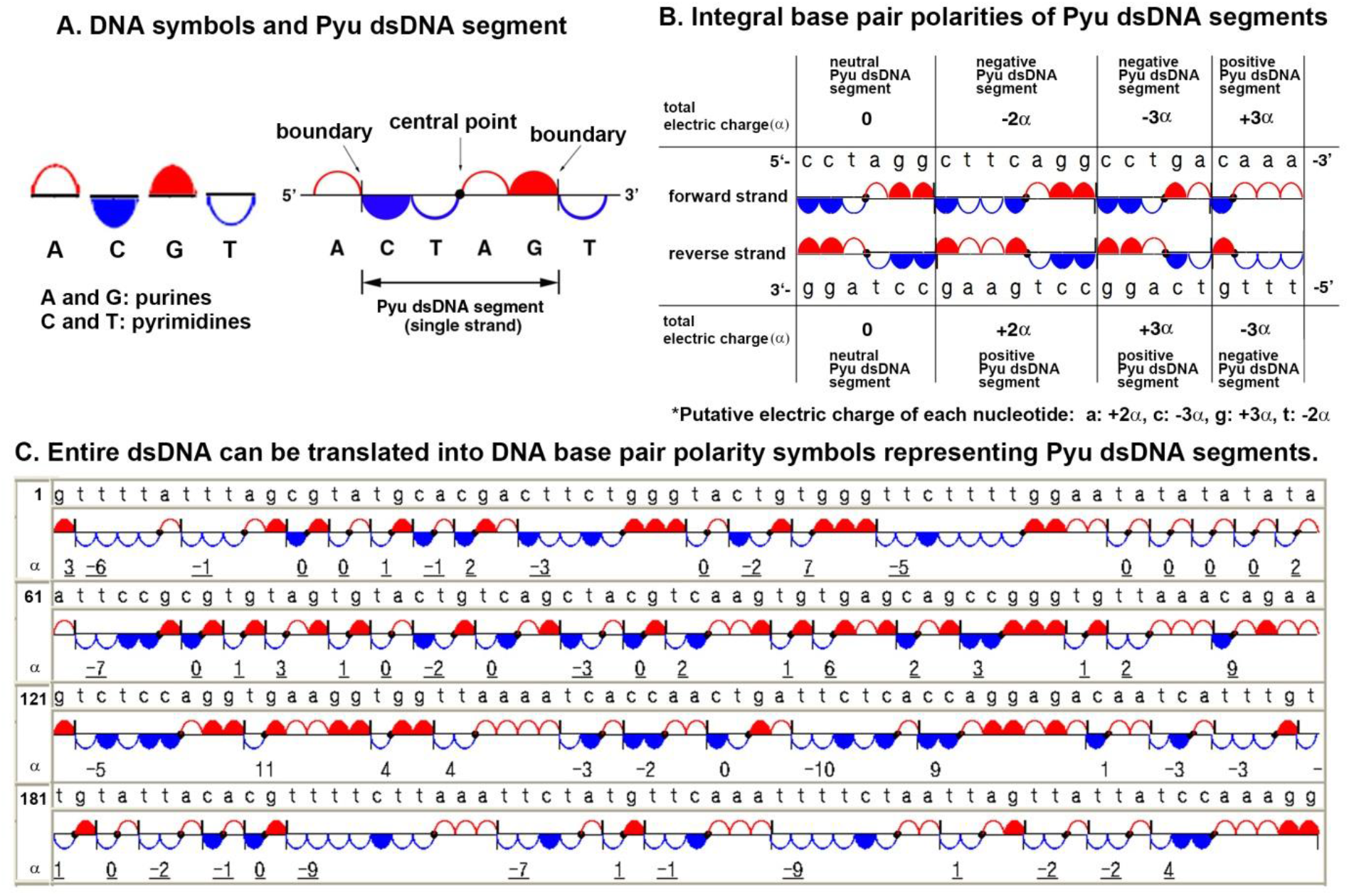
Symbolic illustration of DNA base pair polarities in dsDNA. A: Symbols for A, C, G, and T, and a Pyu segment. B: A Pyu dsDNA consists of two complementary Pyu ssDNAs. C: Whole dsDNA can be divided into Pyu segments.

Hybridization of dsDNA is characteristic and dynamic, depending on the electrostatic hydrogen bonding strength between base pairs, as opposed to base stacking. The increase of hydrogen bonding can induce dominant base pair polarity and rigid conformation of dsDNA, which can interact with different proteins. The present study also showed that Pyu oligo-ssDNA has lower melting temperature (Tm) than Puy oligo-ssDNA, indicating the former hybridizes easily more than the latter (Fig. S1).

We tentatively assigned +2α, -3α, +3α, and -2α base pair polarities to A, C, G, and T, respectively, based on the number of hydrogen bonds in their pairs (where α is a constant value per hydrogen bond) (Fig. 1B).

Therefore, the electrostatic charge of pyrimidines and purines in a Pyu segment can be calculated and combined into α value in the DNA base pair polarity program (*4, 5*), leading the calculation of total DNA hybridization energy (DHE) by Coulomb’s law (Fig. S2).

This study demonstrates that whole dsDNA can be divided into Pyu segments, which appear to be the basic units of genetic code, to preserve the potential energy of dsDNA and to induce optimal DNA conformation for RNA and DNA transcription (Fig. 1C).

### Hybridization potential assay via EtBr intercalation electrophoresis

When the hybridization potential was measured by EtBr intercalation electrophoresis assay, ds12/n(nTnA) (n=1,2,3,4,6,12) were separately prepared by annealing at 70°C and used for electrophoresis with pre-electrophoresis EtBr staining, which can remove loosely bound EtBr during electrophoresis (Supplementary Text 3, Fig. S3). The results showed maximum EtBr fluorescence at ds4(3T3A) and ds3(4T4A), similar to the ensemble enthalpy analyzed by the DINAMelt Server. Whereas the post-electrophoresis staining showed maximum EtBr fluorescence at ds12(TA) and gradually decreased to ds12T12A, which is similar pattern to the total DNA hybridization energy (DHE) analyzed by DNA base pair polarity program (*5*) (Fig. S3, S4).

## Different properties between Pyu and Puy segments

### 1) Pyu oligo-dsDNAs hybridized more than Puy oligo-dsDNAs

The fluorescence of ethidium bromide (EtBr) increases about 25-fold on binding double-stranded DNA (*6*), thereby, EtBr could be used in the hybridization assay to determine the hybridization potential of oligo-dsDNA in 0.1M NaCl solution. The pairs of ss(nTnA) and ss(nAnT) (n=3,4,5,6,7,8,10,12) were purchased (Cosmogenetech Co. Korea) and dissolved in 0.1M NaCl solution at 100 μmol/mL. The samples were heated to 90°C for 5 min and slowly cooled to room temperature to obtain monomeric and dimeric hybridization of ds(nTnA) and ds(nAnT). The dsDNA samples were electrophoresed on 20% polyacrylamide gel (Novex™, USA). EtBr staining was performed before or after electrophoresis to accurately detect EtBr fluorescence under UV light.

Short bps Pyu ds(nTnA) (n=3,4,5,6,7,8) easily underwent self-monomeric hybridization more than Puy ds(nAnT) in post-electrophoresis EtBr staining, and long bps Pyu ds(nTnA) (n=10,12) formed more dimeric hybridization with stronger EtBr staining than Puy ds(nTnA) in both pre- and post-electrophoresis EtBr staining (Fig. 2 A-D). As ds(nTnA) and ds(nAnT) (n=3,4,5,6,7,8,10,12) have simple hybridization of A-T pairing only, it is clear that Pyu ds(nTnA) have stronger monomeric and dimeric hybridization than Puy ds(nAnT) in 0.1 M NaCl solution.

**Fig. 2.**
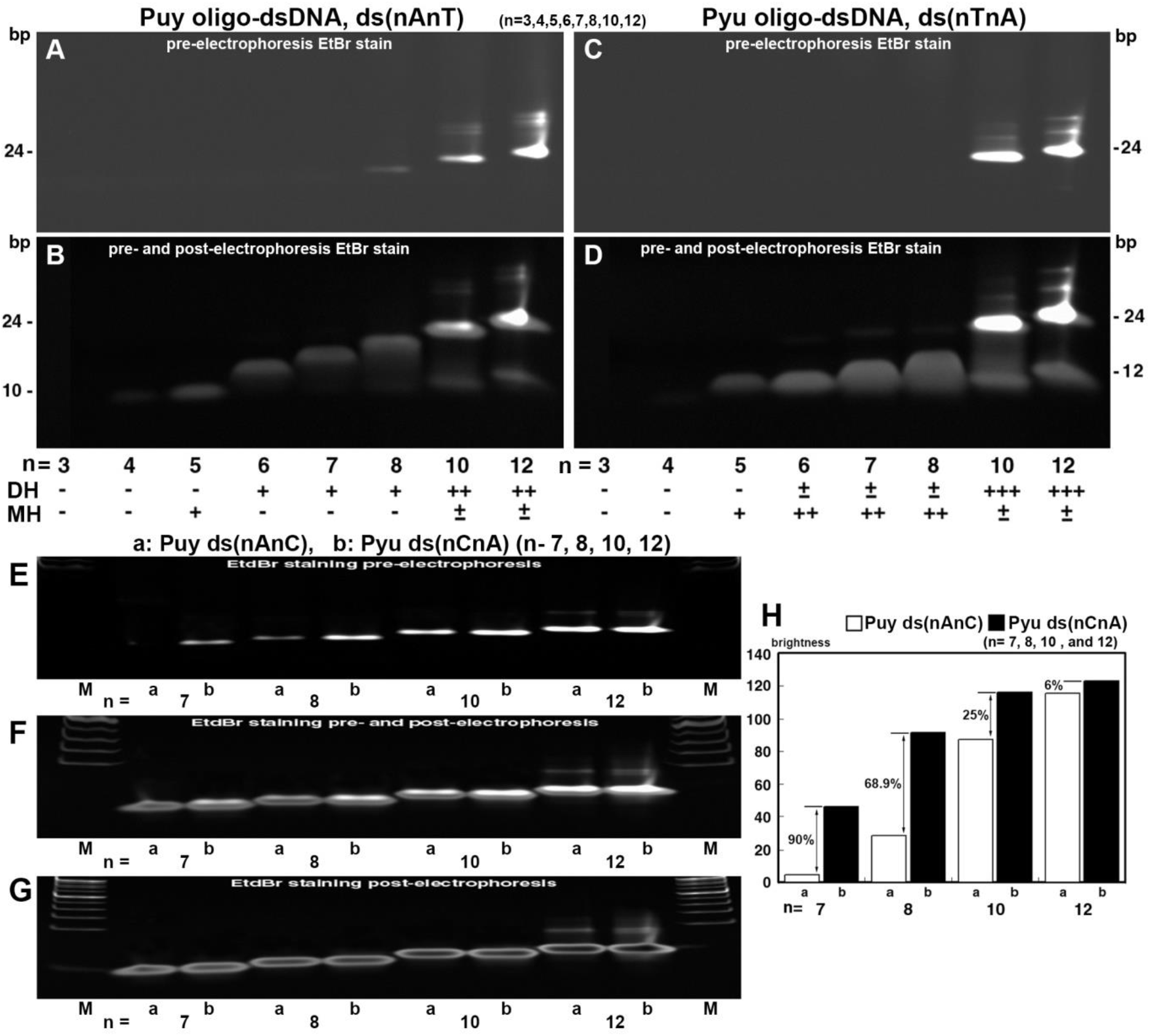
Hybridization status comparing between Puy ds(nAnT) and Pyu ds(nTnA) (n= 3,4,5,6,7,8,10,12) (A-D), Puy ds(nAnC) and Pyu ds(nCnA) (n = 7,8,10,12) (E-H) in pre- and post-electrophoresis EtBr staining. DH: Dimeric hybridization, MH: Self-monomeric hybridization. H: A graph plotted from panel E. This figure represents three repeated experiments.

Since ds(nCnA) and ds(nAnC) (n=7,8,10,12) may show dimeric hybridization only, ds(nCnA) and ds(nAnC) (n=7,8,10,12) in 0.1 M NaCl solution at 100 μmol/mL with complete dimeric hybridization were prepared using the method described above, and analyzed by EtBr intercalation electrophoresis.

As the value of n increased, Pyu ds(nCnA) (n=7,8,10,12) showed significantly higher efficiency in EtBr intercalation compared to Puy ds(nAnC) in pre-electrophoresis EtBr staining (Fig. 2E-G). Specifically, ds7C7A, ds8C8A, ds10C10A, and ds12C12A showed 90%, 68.9%, 25%, and 6% more EtBr intercalation than ds7A7C, ds8A8C, ds10A10C, and ds12A12C, respectively (Fig. 2H). Therefore, it is assumed that Pyu ds(nCnA) (n=7,8,10,12) hybridize more than Puy ds(nAnC) in 0.1M NaCl solution.

### 2) Pyu dsDNAs tended to have larger HPLC peak area than Puy dsDNAs

The conformational differences between Pyu and Puy oligo-dsDNAs in 0.1M NaCl solution could be indirectly assessed by their UV260 absorbance in high performance liquid chromatography (HPLC). The various oligo-dsDNAs in 0.1 M NaCl solution were prepared as described above and analyzed by HPLC using a reverse phase column packed with silica beads and 0.1 M NaCl running buffer at 0.3 mL/min, 30°C.

Among simple dsDNAs consisting of A-T or G-C pairs, Pyu ds3T3A and ds3C3G showed a larger peak area by 4.8% and 29.7% compared to Puy ds3A3T and ds3G3C, respectively. Pyu ds3T3A showed a 144.2% larger peak area than Pyu ds3C3G, and Pyu ds3(TA) showed a 72.8% larger peak area than Pyu ds3(CG). In addition, Pyu ds3(TA) and ds3(CG), consisting of three segments, showed peak areas 70.7% and 141.2% larger than Pyu ds3T3A and ds3C3G, consisting of one segment, respectively (Fig. 3).

**Fig. 3.**
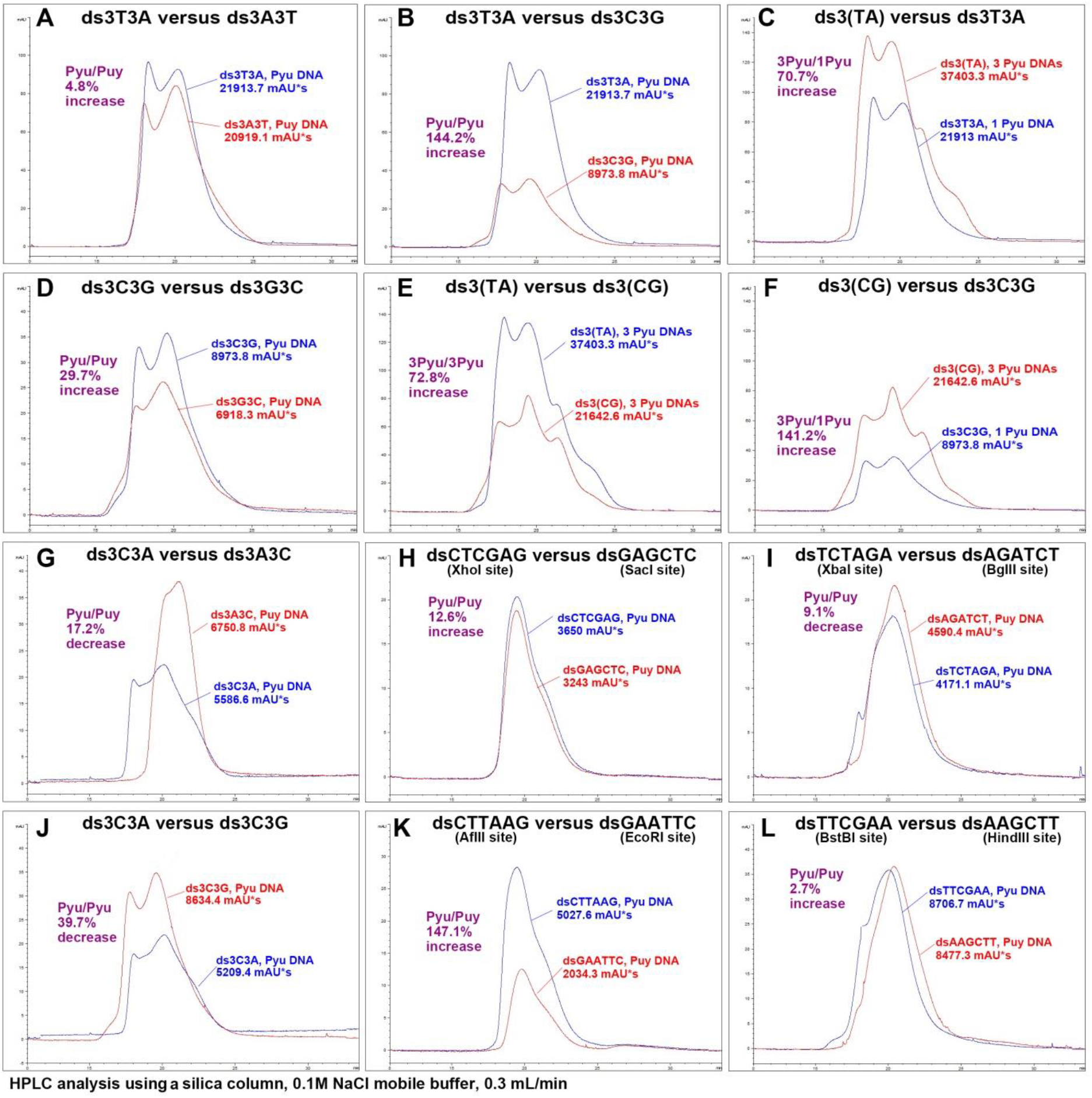
HPLC for simple dsDNA pairs, ds3T3A/ds3A3T (A), ds3C3G/ds3G3C (D), ds3T3A/ds3C3G (B), ds3(TA)/ds3(CG) (E), ds3T3A/ds3(TA) (C), and ds3C3G/ds3(CG) (F), and for complex dsDNA pairs, ds3C3A/ds3A3C (G), ds3C3A/ds3C3G (J), dsCTCGAG/dsGAGCTC (H), dsTCTAGA/dsAGATCT (I), dsCTTAAG/dsGAATTC (K), and dsTTCGAA/dsAAGCTT (L) in 0.1M NaCl solution.

When comparing complex dsDNAs between ds3C3A and ds3A3C, which consist of both A-T and G-C pairs, it was observed that ds3C3A had a faster retention time on reverse phase column but a smaller peak area by 17.2% than ds3A3C. However, ds3C3A had a similar retention time but a 39.7% smaller peak area than ds3C3G (Fig. 3). Based on these results, it can be concluded that complex Pyu ds3C3A is a more condensed form compared to complex Puy ds3A3C and simple Pyu ds3C3G.

When comparing complex dsDNAs of restriction endonuclease (RE) sites, Pyu dsCTCGAG (XhoI) showed a 12.6% larger peak area than Puy dsGAGCTC (SacI). Pyu dsTCTAGA (XbaI) showed a 9.1% larger peak area than Puy dsAGATCT (BgIII). Pyu dsCTTAAG (AfIII) showed a 147.1% larger peak area than Puy dsGAATTC (EcoRI). Pyu dsTTCGAA (BstBI) showed a slightly faster retention time and a 2.7% increase in peak area compared to Puy dsAAGCTT (HindIII) (Fig. 3).

The HPLC peak areas of palindromic dsDNAs encoding different RE sites varied depending on the base pair polarities of Pyu and Puy dsDNAs, although each set of oligo-dsDNAs had the same number of A-T and G-C pairs. There was a tendency for Pyu oligo-dsDNAs to have larger peak area than Puy oligo-dsDNAs. And Pyu oligo-dsDNAs composed of A-T pairs have larger peak areas than those composed of G-C pairs, and Pyu oligo-dsDNAs composed of multiple segments have larger peak areas than those composed of one segment. The results are identical to those in Fig. S1, showing a tendency for Pyu oligo-ssDNAs to have a lower melting temperature (Tm) than Puy oligo-ssDNAs, and for Pyu oligo-ssDNAs composed of A-T pairs to have a lower Tm than those composed of G-C pairs (Supplementary Text 1).

### 3) Pyu oligo-dsDNAs showed higher IR absorbance at 3400-3200 cm^-1^ than Puy oligo-dsDNAs

The pairs of Pyu and Puy oligo-dsDNAs, ds3T3A and ds3A3T, ds3C3G and ds3G3C, ds3C3A and ds3A3C, dsCCCGGG and dsTTAGGG, dsCTCGAG and dsGAGCTC, dsTCTAGA and dsAGATCT were prepared in 0.1M NaCl solution according to the method described above. The N-H stretching of DNA samples were analyzed by FT-IR (Perkin Elmer, USA) by comparing the IR absorbance of O-H stretching in distilled water (DW), 0.1M NaCl, and 1M NaCl solutions.

Pyu ds3T3A, ds3C3G, dsCTCGAG, and dsTCTAGA showed higher IR absorbance at 3400-3200 cm^-1^ than Puy ds3A3T, ds3G3C, dsGAGCTC, and dsAGATCT, respectively, indicating Pyu oligo-dsDNAs have increased N-H stretching in hybridization than Puy oligo-dsDNAs. Conversely, Pyu ds3C3A showed lower IR absorbance than Puy ds3A3C, and Pyu dsTTAGGG, a telomere repeat sequence, showed lower IR absorbance than Pyu dsCCCGGG (Fig. 4). The decrease of IR absorbance in ds3C3A compared to ds3A3C is identical to the decrease of peak area in HPLC conformation assay.

**Fig. 4.**
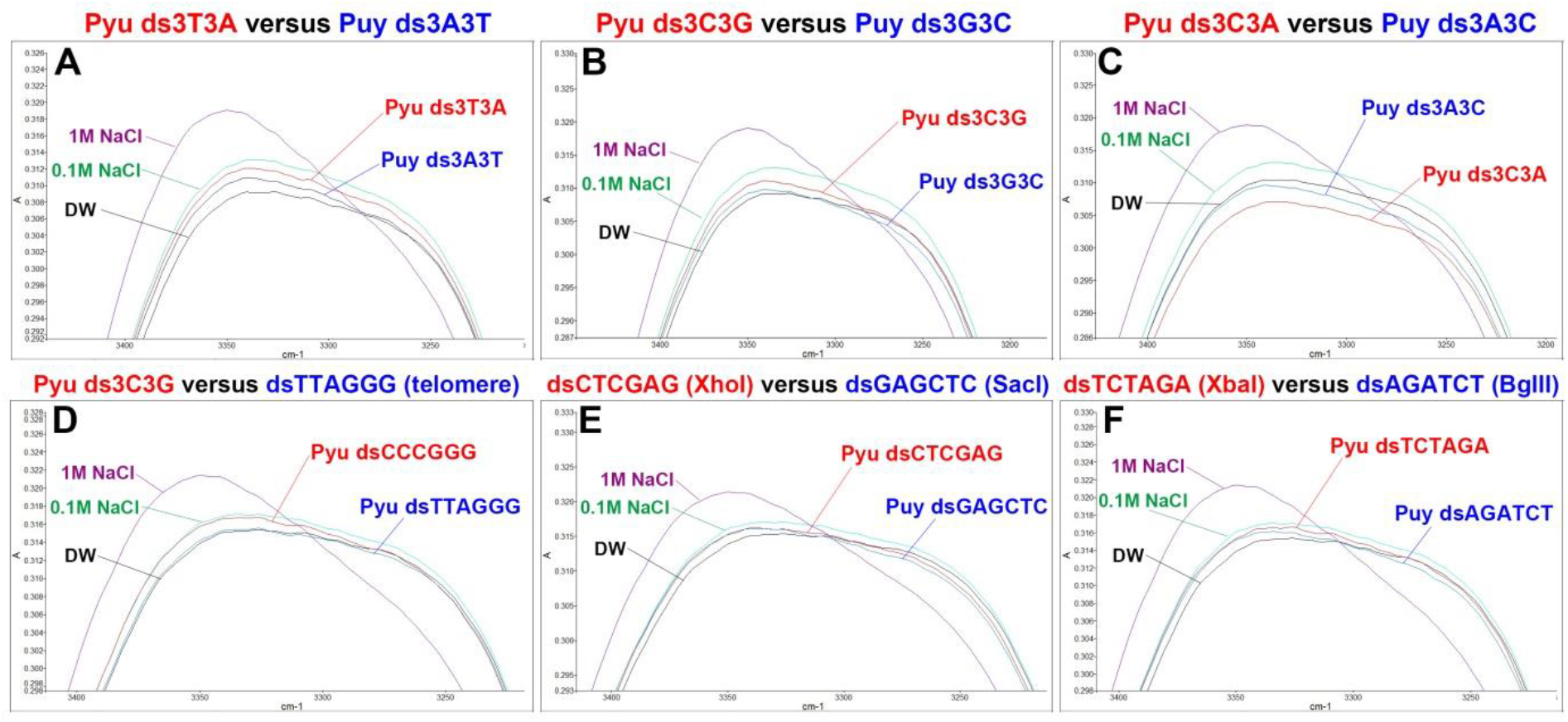
FT-IR for different sets of Pyu and Puy oligo-dsDNA observing IR absorbance at 3400-3200 cm^-1^. The Pyu ds3T3A, ds3C3G, dsCTCGAG, and dsTCTAGA, showed higher IR absorbance than Puy ds3A3T, ds3G3C, dsGAGCTC, and dsAGACTC, respectively.

### 4) In vitro RNA transcription was affected by distinct base pair polarities of inserted oligo-dsDNAs

RNA polymerase may recognize DNA base pair polarities of specific sequences during RNA transcription, and subsequently influence the RNA production. Seven oligo-dsDNAs (approximately 30 bps) with different base pair polarities (Fig. 5D) were separately inserted into a pBluescript vector containing amelogenin cDNA (300 bps) between the SP6 and T7 promoter regions by subcloning with TOPO™ TA cloning kit (Thermo Fisher Scientific Inc. USA). Each subcloned plasmid DNA was linearized by BamHI digestion and used for *in vitro* RNA transcription with SP6 or T7 RNA polymerase (Enzynomics, Korea) at 37°C for 30 min. The RNA products were then immediately electrophoresed on 1.5% formaldehyde agarose gel (20 mM MOPS, 6% formaldehyde) and DEPC-based buffer, stained with EtBr, and detected under UV illumination.

**Fig. 5.**
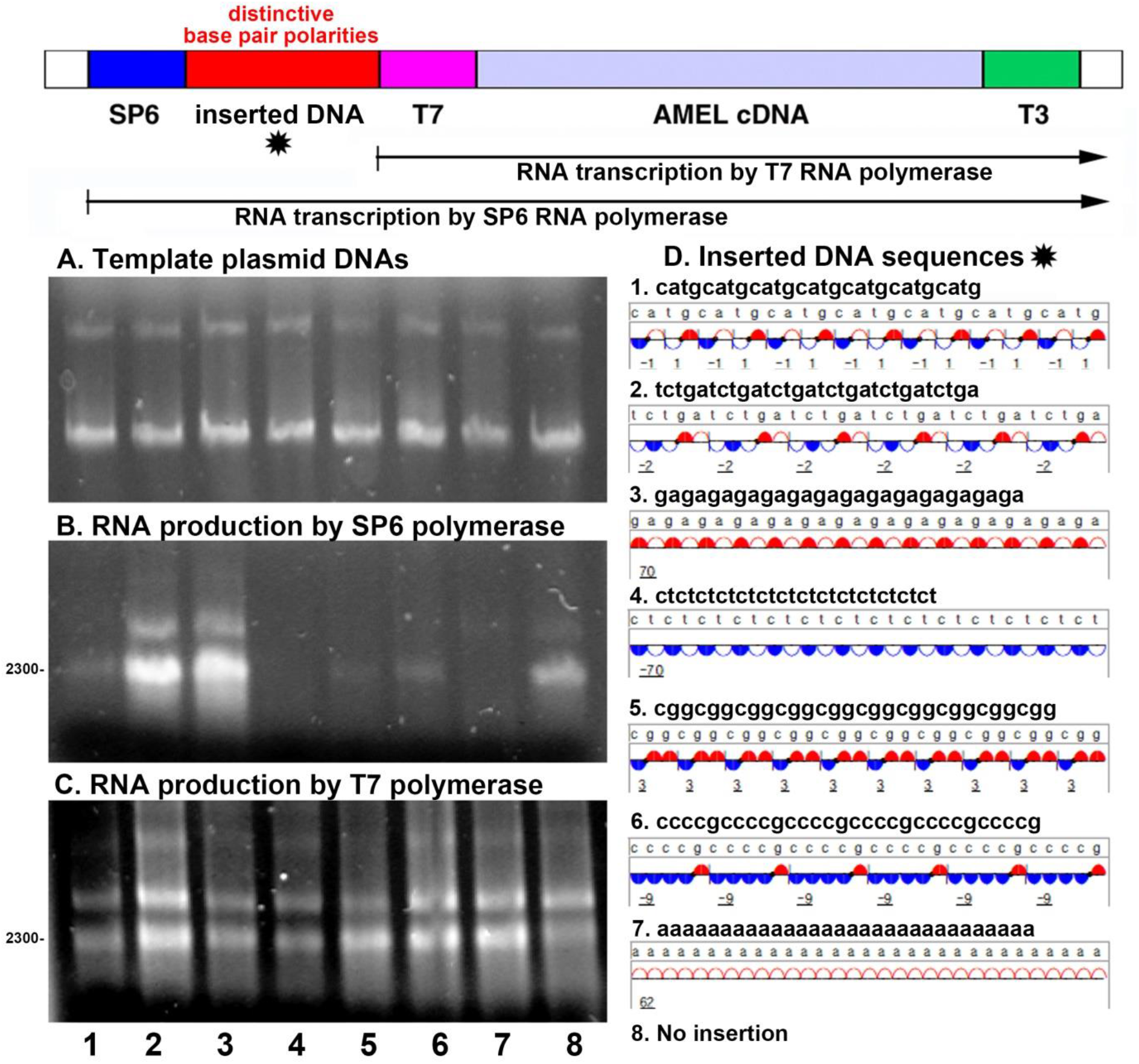
*In vitro* RNA transcription from linear plasmid DNAs inserted with distinct seven oligo-dsDNAs exhibiting different base pair polarities using SP6 or T7 RNA polymerase. A: Linearized template plasmid DNAs. RNA production by SP6 (B) or T7 RNA polymerase (C). D: The inserted DNAs have unique DNA base pair polarities.

*In vitro* RNA transcription resulted in variable RNA production depending on the distinct base pair polarities of inserted DNAs. Compared to the control with no inserted DNA, the template plasmid DNAs inserted with Pyu ds6(TCTGA) and ds14(GA) showed abundant RNA production by SP6 RNA polymerase, while those with ds7(CATG), ds14(CT), ds10(CGG), ds6(CCCCG), and ds30A showed weak RNA production (Fig. 5B). RNA production by T7 RNA polymerase, which is not involved in DNA insertion, appeared to be uniform from all template plasmid DNAs.

The template plasmid DNA inserted with six segments of five bps Pyu dsDNA, ds6(TCTGA), produced more RNA than that with seven segments of four bps Pyu dsDNA forming palindromic pattern, 7(dsCATG). And the template plasmid DNA inserted with 28 bps positive ds14(GA) produced abundant RNA, while those with 28 bps negative ds14(CT) produced much less RNA (Fig. 5B).

On the other hand, the template plasmid DNA inserted with CGG repeats or ds10(CGG), produced a small amount of RNA, and then this RNA production was slightly increased from that with six segments of five bps CG rich Pyu dsDNAs, ds6(CCCCG). While the RNA production from the template plasmid DNA inserted with poly-A sequence, ds30A, was almost inhibited compared to the control from the template plasmid DNA without insertion (Fig. 5B).

### 5) Polymerase chain reaction (PCR) is affected by DNA base pair polarities of primer

Since the primer hybridization on template DNA is critical for PCR, primers of Pyu ds12/n(nCnA) and Puy ds12/n(nAnC) (n=1,2,3,4,6,12) were separately inserted into a multi-cloning site of pBluscript II SK(+) between EcoRI (707) and HindIII (719), and PCR was performed with each primer set, Pyu ss12/n(nCnA) and pBS2, Pyu ss12/n(nTnG) and pBS1, Puy ss12/n(nAnC) and pBS2, and Puy ss(12/n(nGnT) and pBS1 (Fig 6).

**Fig. 6.**
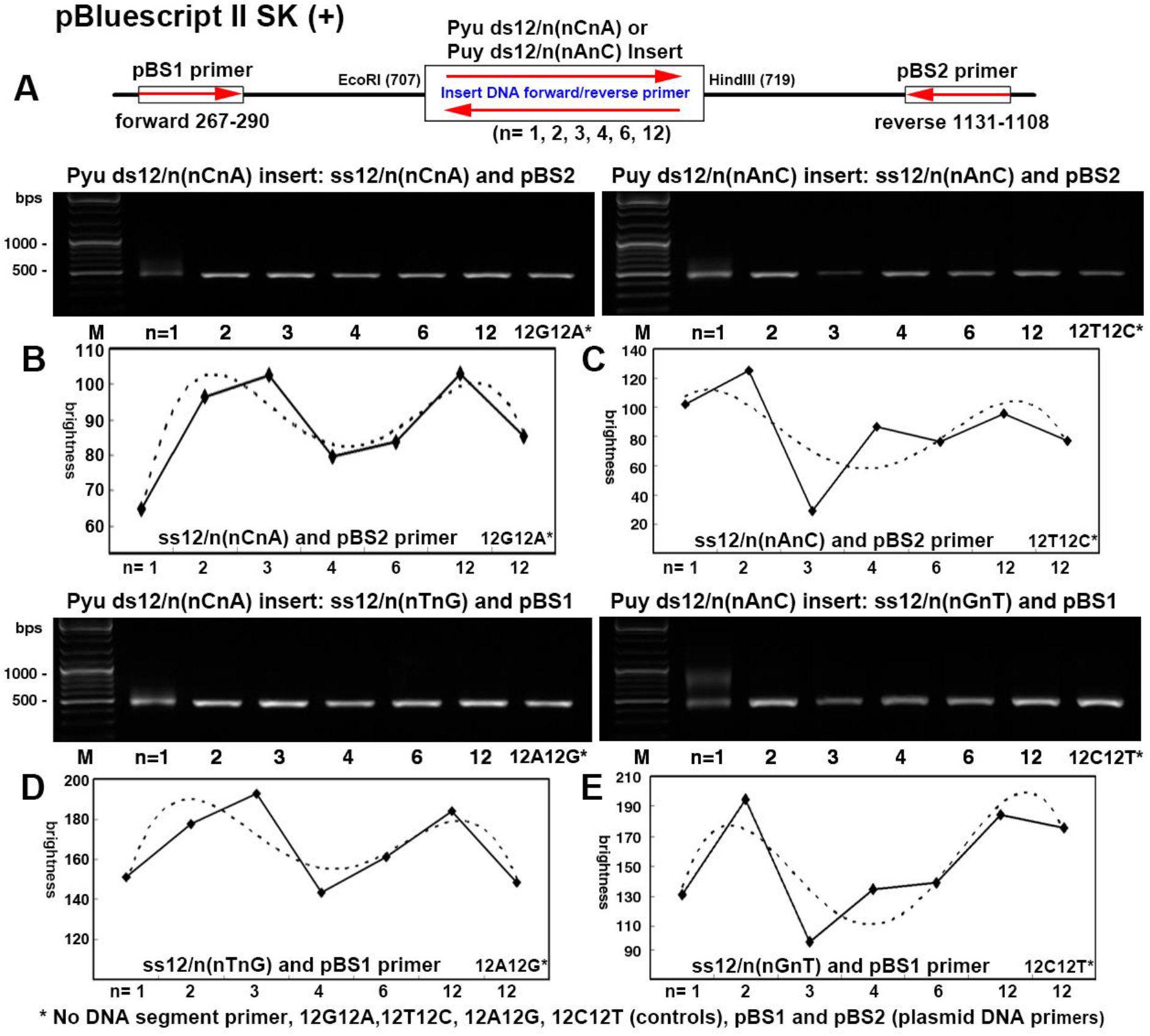
PCR using Pyu or Puy primer on pBluescript II SK (+). Pyu ds12/n(nCnA) and Puy ds12/n(nAnC) were separately inserted into plasmid DNA between EcoRI (707) and HindIII (719) sites. Pyu primers, ss12/n(nCnA) and ss12/n(nTnG) produced more DNA than Puy primers, ss12/n(nAnC) and ss12/n(nGnT).

The results showed that Pyu primers produced stronger bands than Puy primers, and the maximum PCR product was found when using Pyu primers consisting of large segments, 4(3C3A) and 4(3T3G), while when using Puy primers consisting of small segments, 6(2A2C) and 6(2G2T) (Fig. S5), indicating that Pyu primers can easily hybridize on template DNA and subsequently produce abundant DNA compared to Puy primers.

### 6) PCR primers analyzed by DNA base pair polarity

PCR using different primer sets from pBluescript II SK (+), which show different base pair polarities (Fig. 7A). The primer set consisting of multiple segments with complicated polarities (6, 8, 11) produced abundant DNA compared to those consisting a few segments with monotonous polarities (1, 2, 4). The palindromic sequence may inhibit the PCR (3, 5, 6, 7, 8, 9, 10) due to primer dimerization. Particularly, the presence of palindromic sequence at 3’ end of primer (5, 7, 9, 10) greatly affect the PCR production (Fig. 7B).

**Fig. 7.**
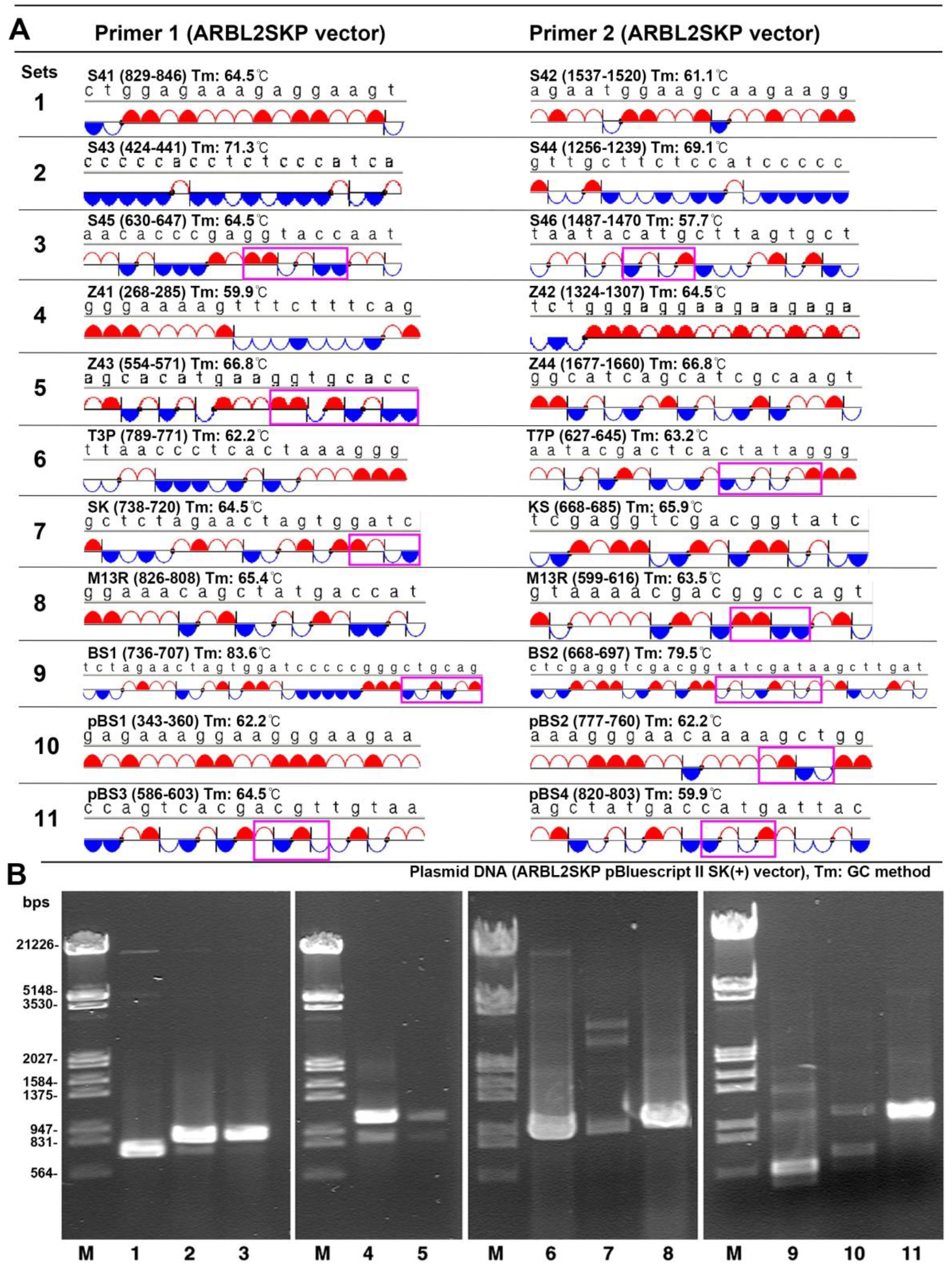
PCR using different primer sets from pBluescript II SK (+), which show different base pair polarities (A). B: The primer set consisting of multiple segments with complicated polarities (6, 8, 11) produced abundant DNA compared to those consisting a few segments with monotonous polarities (1, 2, 4). The palindromic sequence may inhibit the PCR (3, 5, 6, 7, 8, 9, 10) due to primer dimerization. Particularly, the presence of palindromic sequence at 3’ end of primer (5, 7, 9) greatly affect the PCR production. Pink square: palindromic sequences.

## Discussion

In this study, DNA base pair polarities could be illustrated by simple DNA symbols and Pyu/Puy dsDNA segments, which differentially affect the hybridization potential and DNA conformation in a sequence-specific manner. Pyu oligo-dsDNAs tended to have higher hybridization potential and unique conformation, and showed increased IR absorbance than Puy oligo-dsDNAs. *In vitro* RNA transcription was enhanced by the insertion of six segments of five bps Pyu dsDNA (ds6(CTGAG)) and positive ds14(GA), whereas it was inhibited by seven segments of four bps Pyu dsDNA repeats (7(dsCATG)) and negative ds14(CT) compared to the control. On the other hand, G-C pair rich ds10(CGG), ds6(CCCCG) and poly-A (ds30A), which may have tertiary structure, strongly inhibited *in vitro* RNA transcription. In PCR assay, it appears that Pyu primers can easily hybridize on template DNA and subsequently produce abundant DNA compared to Puy primers

Actually, many DNA motif sequences have specific Pyu segment(s) as a core element, i.e., Kozac consensus sequence: dsG(CCG)(CCR)(CCA)TG, telomere repeat sequence: ds(TTAGGG), canonical E-box sequence: ds(CA)(CG)(TG), GLI binding site sequence: dsGA(CCA)(CCCA), HMG1 binding site sequence: ds(TTCA)(TTCA)(TTCA), NF1 binding site sequence: ds(CCAAT), TATA box sequence: ds(TA)(TA)(TAA), SMAD binding site sequence: ds(TCTAGA), RUNX binding site sequence: dsA(CCR)(CCA), etc., ((): A Pyu segment, R: purine, A or G). These Pyu segments may be readily activated with their unique base pair polarities and conformations, and subsequently exert the functions of DNA motifs.

Therefore, it is postulated that Pyu dsDNA could be a basic unit of genetic code to preserve DNA signals and support various DNA functions, and that the semiconductive Pyu and Puy dsDNAs, which are able to preserve and accumulate their unique base pair polarities as an electrostatic charge, may be responsive to a specific external electromagnetic wave. However, this phenomenon remains to be elucidated in the following studies.

However, it is obvious that the dsDNAs, which are composed of only two kinds of pair, A-T and G-C, but are capable of forming complicated coiled coil and tertiary structures with their molecular polarities, must define their molecular characteristics in some simple way available for computational analysis. This study proposes a concept of DNA base pair polarity that may have implications for DNA structure and function.

## Acknowledgments

We would like to express our gratitude to the late Professor Je Geun Chi and the late Dr. Soo Il Chung, who contributed to this research in part.

