## Supplement Fig S1-S4 for "DNA Base Pair Polarities of Pyrimidine(s)-Purine(s) (Pyu) dsDNA induce stronger Hybridization and unique DNA Conformation compared to Purine(s)-Pyrimidine(s) (Puy) dsDNA"

**The PDF file includes:**

Materials and Methods
Supplementary Text
Figs. S1 to S4
References

### Materials and Methods

#### *1) Comparison of DNA melting temperature (Tm) between simple Puy and Pyu ssDNA*

Tm values of Pyu ss6/n(nTnA), Pyu ss6/n(nCnG), Puy ss6/n(nAnT), and Puy ss6/n(nGnC) (n=1,2,3,6), which consist of A-T or G-C pairs only, were measured via Kun's Oligonucleotide Tm calculator website (Nearest-Neighbor method, Harvard University) (*1*, *2*). The results were plotted in a graph to compare Tm values between Pyu and Puy oligo-ssDNAs.

#### *2) Hybridization of oligo-dsDNAs*

To prepare the palindromic sequence oligo-dsDNAs, including as ds(nTnA) and ds(nAnT) (n=2,4,5,6,7,8,9,10,12), ds12/n(nTnA) and ds12/n(nAnT) (n=1,2,3,4,6,12), ds3T3A, ds3A3T, ds3C3G, ds3G3C, etc., each oligo-ssDNA was purchased from Cosmogenetech Co. Ltd. (Korea, Seoul) and dissolved in 0.1M NaCl solution at 200 μmol/mL. The samples were heated to 90°C for 10 minutes and gradually cooled to room temperature and then each 100 μmol/mL oligo-dsDNA in 0.1M NaCl solution was obtained and immediately used for the following experiment.

To prepare the complicated sequence oligo-dsDNAs, including ds3C3A, ds3A3C, ds12/n(nCnA) and ds12/n(nAnC) (n=1,2,3,4,6,12), dsTTAGGG, etc., each pair of oligo-dsDNA was purchased from Cosmogenetech Co. Ltd. (Korea, Seoul) and dissolved in 0.1M NaCl solution at 100 μmol/mL. The samples were heated to 90°C for 10 minutes and gradually cooled to room temperature, and then 100 μmol/mL oligo-dsDNA in 0.1M NaCl solution was obtained and immediately used for the following experiment.

#### *3) Electrophoresis for oligo-dsDNAs and plasmid DNAs*

The small size oligo-dsDNAs less than 500 bps were electrophoresed on 20% polyacrylamide gel (Novex™, USA) using TBE buffer, while the large size DNA products from PCR, RE digestion, *in situ* RNA transcription on plasmid DNA were electrophoresed on 1-2% agarose gels using TAE buffer.

Gels were stained with EtBr before or after electrophoresis. The pre-electrophoresis EtBr stain may show a distinct DNA band indicating potential EtBr intercalation by removing the incomplete EtBr binding during electrophoresis, while the post-electrophoresis EtBr stain may show the total amount of EtBr bound to DNA.

#### *4) HPLC analysis for DNA conformationalchanges*

Since the UV260 absorbance in DNA molecules is variable depending on the structures of DNA conformation, HPLC at UV260 is available to detect the conformational changes of oligo-dsDNA in 0.1M NaCl solution. This study performed HPLC using a reverse phase column packed with non-adherent silica beads (Daisogel SP-300-5P, Osaka Soda Co., Ltd., Japan) at low speed mobile phase, 0.3-0.5 mL/min, with 0.1M NaCl solution, 30℃. The precision HPLC (1100, Agilent, USA) was equipped with UV spectroscope (DAD), fluoroscope (FLD), and auto-sampler.

#### *5) FT-IR analysis for oligo-ds DNAs*

The pairs of Pyu and Puy oligo-dsDNAs, ds3T3A and ds3A3T, ds3C3G and ds3G3C, ds3C3A and ds3A3C, dsCCCGGG and dsTTAGGG, dsCTCGAG and dsGAGCTC, dsTCTAGA and dsAGATCT were prepared in 0.1M NaCl solution according to the method described above.

The N-H stretching of DNA samples were analyzed by FT-IR (Perkin Elmer, USA) by comparing the IR absorbance of O-H stretching in distilled water (DW), 0.1M NaCl and 1M NaCl solutions.

***6) In vitro RNA transcription assay for DNA base pair polarity***

RNA polymerase can recognize the DNA base pair polarities of specific sequences during RNA transcription, and subsequently influence the RNA production. Seven oligo-dsDNAs (approximately 30 bps) with different base pair polarities (Fig. 5 D) were separately inserted into a pBluescript vector containing amelogenin cDNA (300 bps) between the SP6 and T7 promoter regions by subcloning with TOPO™ TA cloning kit (Thermo Fisher Scientific Inc. USA). Each subcloned plasmid DNA was linearized by BamHI digestion and used for *in vitro* RNA transcription with SP6 or T7 RNA polymerase (Enzynomics, Korea) at 37°C for 30 min. The RNA products were then immediately electrophoresed on a 1.5% formaldehyde agarose gel (20 mM MOPS, 6% formaldehyde) and DEPC-based buffer, stained with EtBr, and detected under UV illumination.

***7) Polymerase chain reaction (PCR) for DNA base pair polarity***

To determine the difference between Pyu and Puy primers in PCR, Pyu ds12/n(nCnA) and Puy ds12/n(nAnC) (n=1,2,3,4,6,12) were separately inserted into a multi-cloning site of pBluscript II SK(+) between EcoRI (707) and HindIII (719) by subcloning with TOPO™ TA cloning kit (Thermo Fisher Scientific Inc. USA). PCR was performed using the subcloned plasmid DNA between the inserted DNA primers, Pyu ss12/n(nCnA), Pyu ss12/n(nTnG), Puy ss12/n(nAnC), Puy ss12/n(nGnT) and plasmid primers pBS1

(TGCCGTAAAGCACTAAATCGGAACC (267-290)) and pBS2 (ATTCTGTGGATAACCGTATTACCG (1131-1108)).

### Supplementary Text

#### *1) Comparison of Tm value between Puy and Pyu ssDNAs*

In this study, it was found that Pyu ss6/n(nTnA) and Pyu ss6/n(nCnG) (n=1,2,3,6) have lower melting temperature (Tm) than Puy ss6/n(nAnT) and Puy ss6/n(nGnC), indicating the formers hybridize more easily than the latters (Fig. S1).

Particularly, ss6(AT) and ss6(TA) which have 6 segments showed much lower Tm than ss6A6T and ss6T6A which have one segment, while ss6(GC) and ss6(CG) which have 6 segments showed much higher Tm than ss6G6C and ss6C6G which have one segment. The results indicate Tm of ss6/n(nTnA) and ss6/n(nAnT) consisting A-T pairs is contrary to Tm of ss6/n(nCnG) ss6/n(nGnC) (n=1,2,3,6), respectively (Fig. S1).

Therefore, it is postulated that among oligo-dsDNAs of the same length, simple oligo-ssDNA consisting of A-T pairs become regular and monotonous structure as the number of segment increases, while simple oligo-ssDNAs consisting of G-C pairs become irregular and complicate structures which are difficult to hybridize.

#### *2) Diagram of DNA base pair polarity program*

A concept of DNA base pair polarity is based on the interaction between hybrid molecules that have different molecular polarities from each other, resulting in the formation of a complementary double-stranded DNA (dsDNA). The dsDNA may not show intramolecular electron transfer due to the presence of multiple carbon-carbon bonds (sigma bonds) in its axis, and resulted in a dielectric molecule.

However, dsDNA is electrostatically polarized by the negative phosphate group and positive base residue. When two bases are hybridized by hydrogen bonding, the bases are electrostatically polarized positively or negatively depending on the dipole hydrogen bond. Here, purine is positive due to the presence of cationic imidazole ring compared to pyrimidine, and two hybrid bases reach equilibrium by hydrogen bonding and then are electrostatically replaced by distant phosphate group.

This unique molecular polarity of dsDNA can be symbolized and summarized into a hypothetical concept of DNA base pair polarity, which leads the characterization of Pyu (pyrimidine(s)-purine(s)) dsDNA segments. The electrostatic charge of pyrimidines and purines in a Pyu segment can be analyzed by Coulomb's law (*3-5*) by combining them into α value in the DNA base pair polarity program  to calculate total DNA hybridization energy (DHE) (*6*, *7*) (Fig. S2).

This unique molecular polarity of dsDNA can be symbolized and summarized in a hypothetical concept of DNA base pair polarity, which leads to the characterization of Pyu (pyrimidine(s)-purine(s)) dsDNA segments. The electrostatic charge of pyrimidines and purines in a Pyu segment can be analyzed by Coulomb's law (3-5) by combining them into the α value in the DNA base pair polarity program to calculate the total DNA hybridization energy (DHE) (6, 7) (Fig. S2).

***3) Hybridization potential assay via EtBr intercalation electrophoresis***

Since DNA hybridization is greatly affected by annealing temperature, the hybridization potential can be hindered by low annealing temperature below the melting temperature (Tm), while denatured and complete hybridization can be achieved by high annealing temperature above 90℃. Therefore, to know the hybridization potential of ds12/n(nTnA) (n=1,2,3,4,6,12), the optimal annealing temperature should be determined first.

The hybridization assay of ds12/n(nTnA) was performed by hybridizing ss12/n(nTnA) separately at 20℃, 37.5℃, 70℃, and 90℃, because the Tm of ss12/n(nTnA) ranges from 39.36℃ to 47.79℃. After annealing for 10 min, each ds12/n(nTnA) was slowly cooled to room temperature and then electrophoresed on 20% polyacrylamide gel (Novex™, USA) with EtBr staining before and after electrophoresis. EtBr fluorescence in the gel was observed under UV light.

The hybridization of ds12/n(nTnA) at the annealing temperature of 20℃, 37.5℃, 70℃, and 90℃ showed that ds12/n(nTnA) was incompletely hybridized by the annealing temperature of 20℃ and 37.5℃, while completely hybridized by 90℃, hardly showing the difference of hybridization potential between ds12/n(nTnA). While the annealing temperature of 70℃ resulted in distinguishable hybridization potential between ds12/n(nTnA). Therefore, the hybridization results at 70℃ were analyzed to determine the hybridization potential depending on the number of segments as shown in Fig. 4S.

Palindromic ss12/n(nTnA) (n=1,2,3,4,6,12) were separately dissolved in 0.1M NaCl solution and heated to 70 ℃ for 10 min and slowly cooled to room temperature. The samples were electrophoresed on 20% polyacrylamide gel (Novex™, USA) with EtBr staining before or after electrophoresis and visualized under UV light.

Pre-electrophoresis EtBr staining showed clear bands, while post-electrophoresis EtBr staining showed blurred bands. The bands by dimeric hybridization in pre-electrophoresis EtBr staining showed dominant brightness in ds4(3T3A) and ds3(4T4A), which was similar pattern to the increased ensemble enthalpy measured via DINAMelt server (Rensselaer Polytechnic Institute).

On the other hand, the bands of both dimeric and monomeric hybridization in post-electrophoresis EtBr staining showed the maximum brightness in ds12(TA), but gradually decreased brightness and resulted in the minimum brightness in ds12T12A, which was similar pattern to the total DNA hybridization energy (DHE) calculated via DNA base pair polarity program in this study. The results indicate the DHE may be identical to the amount of total EtBr binding to oligo-dsDNA in post-electrophoresis staining.

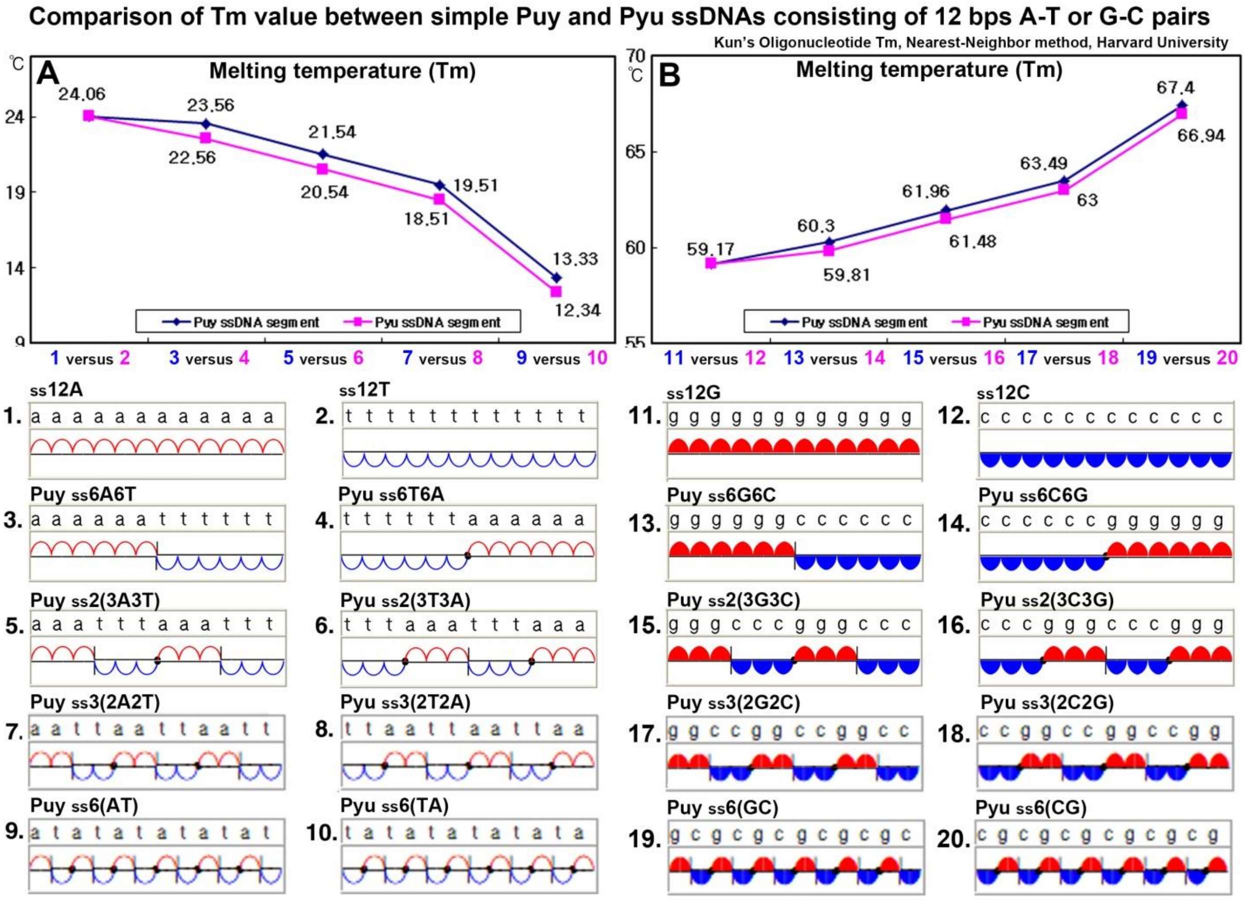


**Fig. S1.** Comparison of DNA melting temperature (Tm) between simple Puy and Pyu ssDNA consisting of 12 bps A-T or G-C pairs. Simple Puy and Pyu ssDNAs consisting of A-T pairs (A) or G-C pairs (B) exhibiting distinctive ssDNA polarity and different number of segment show contrasting Tm values.

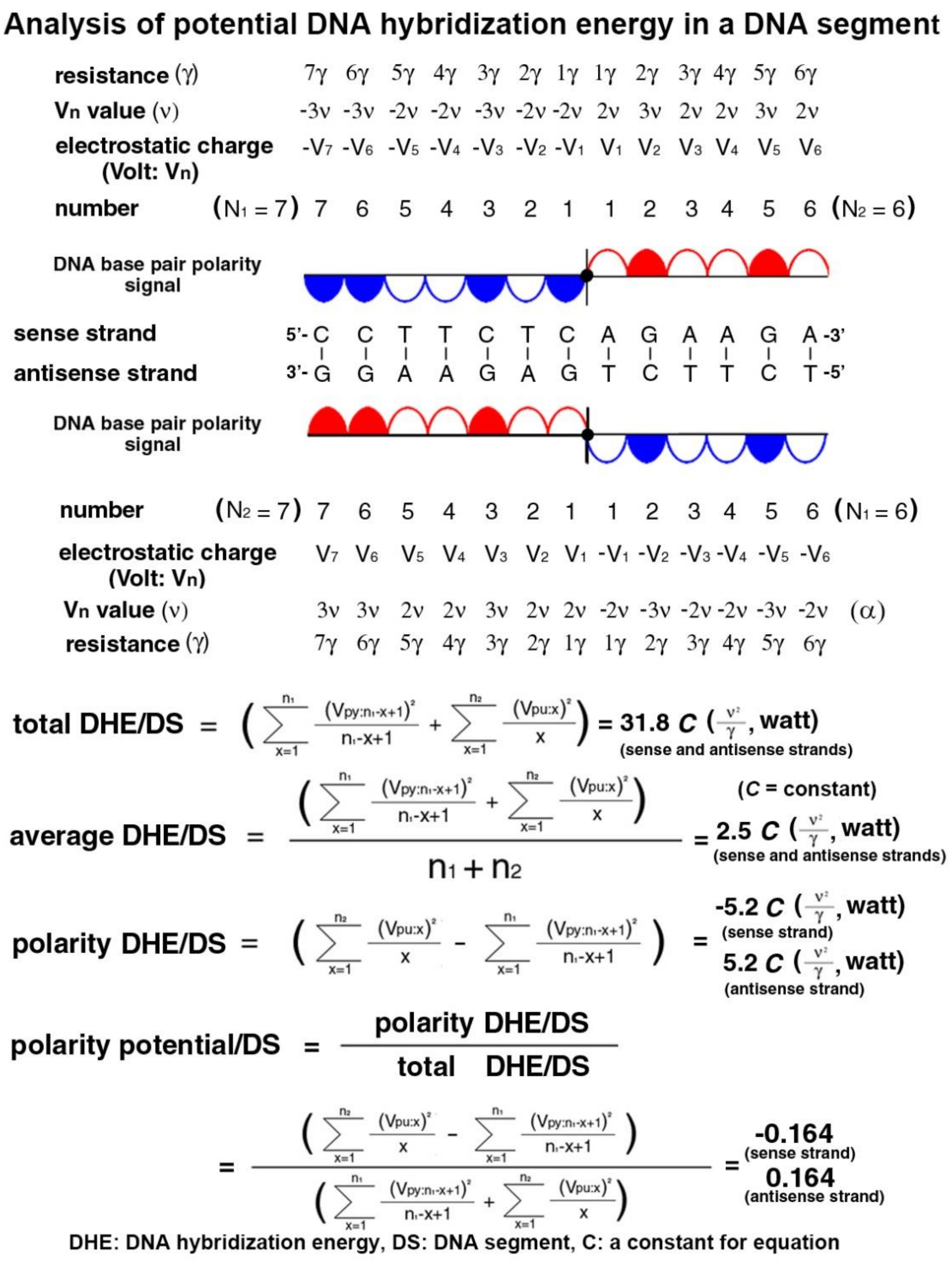


**Fig. S2.** Diagram of DNA base pair polarity program. The electrical charge in a dsDNA segment could be calculated to get total DNA hybridization energy (DHE) via Coulomb's law.

D

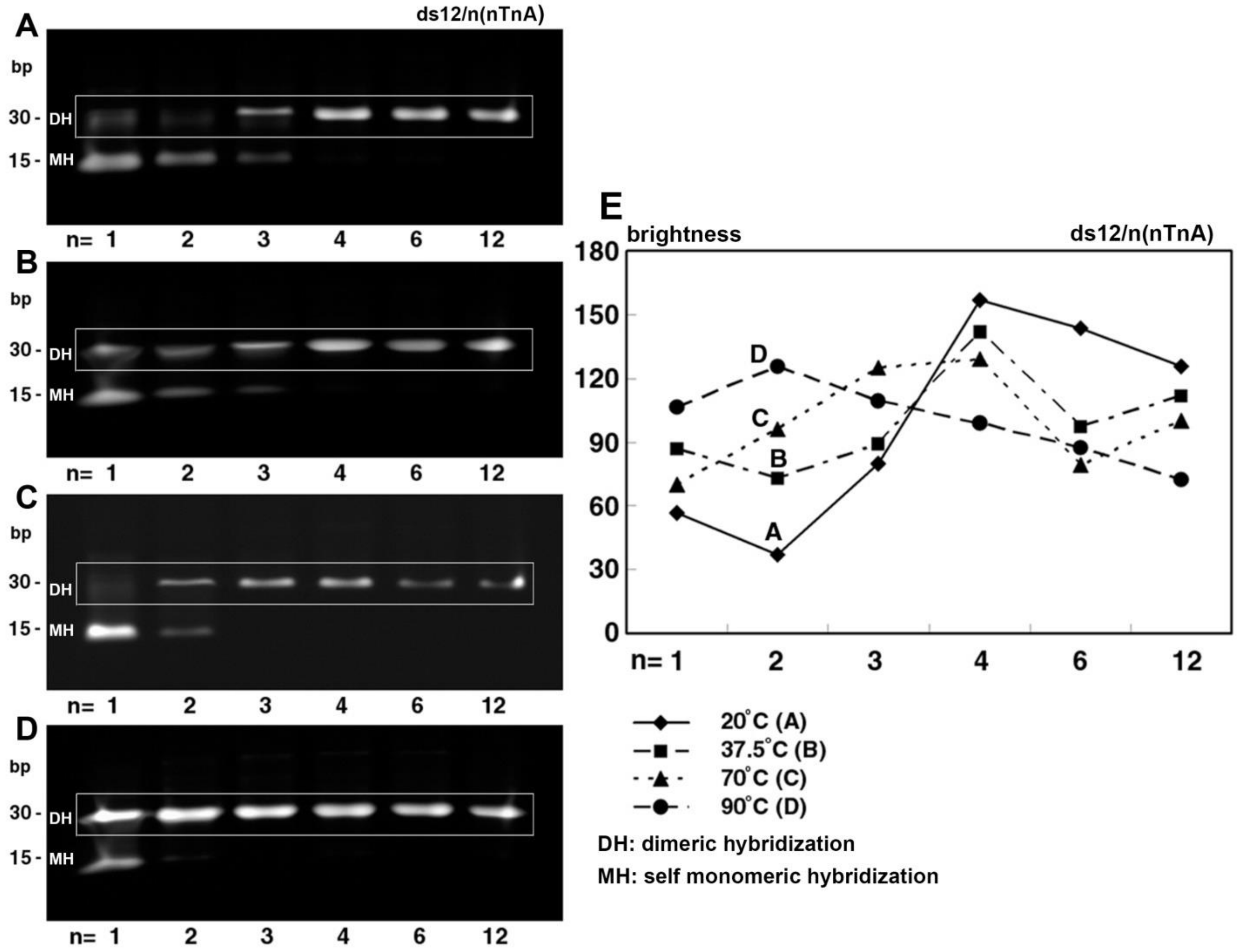


**Fig. S3**. DNA hybridization to determine the optimal annealing temperature for the hybridization potential assay of ds12/n(nTnA) (n=1,2,3,4,6,12). Each palindromic ss12/n(nTnA) was annealed at 20℃ (A), 37.5℃ (B), 70℃ (C), and 90℃ (D). E: A graph plotted with all data.

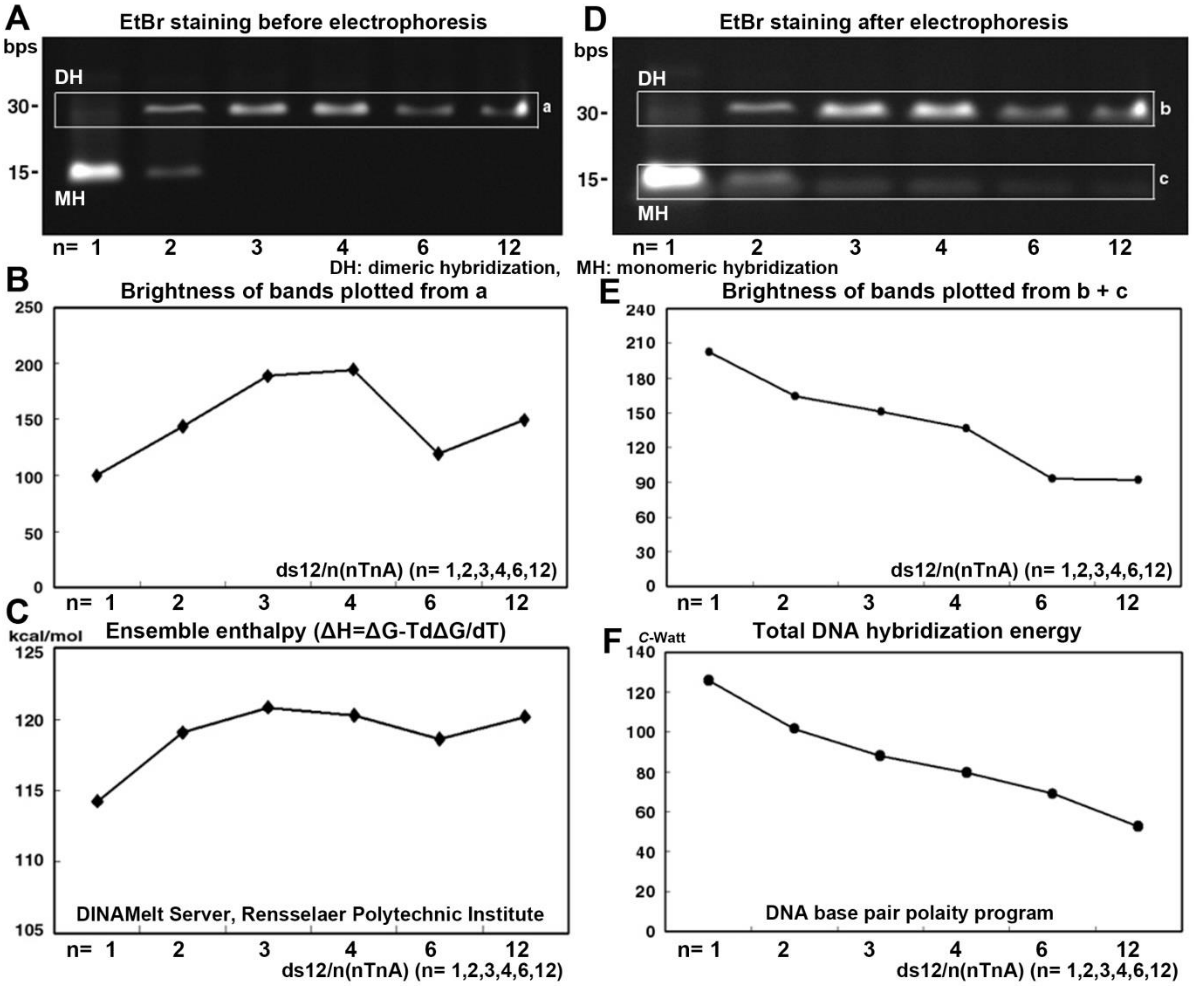


**Fig. S4.** Hybridization potential assay for ds12/n(nTnA) (n=1,2,3,4,6,12) annealed at 70 ℃. A-C: Bands by dimeric hybridization were analyzed for EtBr brightness and ensemble enthalpy (DINAMelt server). D-F: Bands by both of dimeric and monomeric hybridization were analyzed for EtBr brightness and total DNA hybridization energy (DNA base pair polarity program in this study)
